# Tasting Pictures: Viewing Images of Foods Evokes Taste-Quality-Specific Activity in Gustatory Insular Cortex

**DOI:** 10.1101/2020.10.14.307454

**Authors:** Jason A. Avery, Alexander G. Liu, John E. Ingeholm, Stephen J. Gotts, Alex Martin

## Abstract

Previous studies have shown that the conceptual representation of food involves brain regions associated with taste perception. The specificity of this response, however, is unknown. Does viewing pictures of food produce a general, non-specific response in taste-sensitive regions of the brain? Or, is the response specific for how a particular food tastes? Building on recent findings that specific tastes can be decoded from taste-sensitive regions of insular cortex, we asked whether viewing pictures of foods associated with a specific taste (e.g., sweet, salty, sour) can also be decoded from these same regions and if so, are the patterns of neural activity elicited by the pictures and their associated tastes similar? Using ultra-high resolution functional magnetic resonance imaging at high magnetic field strength (7-Tesla), we were able to decode specific tastes delivered during scanning, as well as the specific taste category associated with food pictures within the dorsal mid-insula, a primary taste responsive region of brain. Thus, merely viewing food pictures triggers an automatic retrieval of specific taste quality information associated with the depicted foods, within gustatory cortex. However, the patterns of activity elicited by pictures and their associated tastes were unrelated, thus suggesting a clear neural distinction between inferred and directly experienced sensory events. These data show how higher-order inferences derived from stimuli in one modality (i.e. vision) can be represented in brain regions typically thought to represent only low-level information about a different modality (i.e. taste).

**Significance Statement:** Does a picture of an apple taste sweet? Previous studies have shown that viewing food pictures activates brain regions involved in taste perception. However, it’s unclear if this response is actually specific to the taste of depicted foods. Using ultra-high resolution functional magnetic resonance imaging and multi-voxel pattern analysis, we decoded specific tastes delivered during scanning, as well as the dominant tastes associated with food pictures within primary taste cortex. Thus, merely viewing pictures of food evokes an automatic retrieval of information about the taste of those foods. These results show how higher-order information from one sensory modality (i.e. vision) can be represented in brain regions thought to represent only low-level information from a different modality (i.e. taste).

## Introduction

Throughout the course of their lives, individuals have endless experiences with different types of food. Through these experiences, we grow to learn the predictable associations between how foods look, smell, and taste, as well as how nourishing they are. Based upon numerous individual examples, we learn that ice cream tastes sweet, pretzels taste salty, and lemons taste sour. These sight-taste associations allow us to form a richly detailed conceptual model of the foods we experience, which we can utilize to predict how a novel instance of this food will taste. The continued popularity and utility of visual advertisements for driving food sales attests to the power of these associations to motivate consumptive behavior.

Grounded theories of cognition, supported by decades of human neuroimaging evidence, claim that object concepts are represented, in part, within the neural substrates involved in perceiving and interacting with those objects(1, 2). Within this view, the conceptual representation of food should likewise involve the brain regions associated with taste perception and reward. This possibility has been borne out by human neuroimaging studies that have shown that viewing food pictures elicits activity in a taste-responsive brain regions such as the insula, amygdala, and orbitofrontal cortex (OFC)(3–6) (for meta-analyses, see (7, 8)). Within these studies, activation of the bilateral mid-insular cortex when viewing food pictures is of particular interest because of its central role in taste perception(4, 9–12) and putative role as human primary gustatory cortex(13–15). Indeed, there is some evidence that viewing food pictures can even modulate taste-evoked neural responses within the mid-insular cortex(16). Taken together, these studies suggest that viewing food pictures triggers the automatic and implicit retrieval of taste property information from gustatory cortex. However, it is unclear whether these insular activations to food pictures represent a general taste-related response, or whether they represent the retrieval of specific taste property information, such as whether a food tastes predominantly sweet, salty, or sour, as no study has investigated whether food pictures activate gustatory cortex in a taste-quality-specific manner.

Indeed, this question might prove nearly intractable using standard neuroimaging analyses and relatively low-resolution functional imaging methods. Recent studies using multivariate pattern analysis (MVPA) applied to high-resolution fMRI data, however, suggest that taste quality is represented by distributed patterns of activation within taste-responsive regions of the brain, rather than topographically(12, 17, 18). These findings raised the possibility that inferred taste qualities evoked by viewing food pictures might also be discernable in gustatory cortex using a similar approach. If this is indeed the case, an important and related question would be, are the distributed activity patterns used to represent the inferred taste qualities associated with food pictures the same as, or reliably similar to, the activation patterns which represent experienced tastes?

In order to address these questions, we conducted an fMRI study in which we had subjects undergo high resolution 7-Tesla fMRI while performing tasks in which they viewed pictures of foods which varied in their predominant taste quality (sweet, salty, or sour), as well as pictures of non-food objects. During the same scan session, participants performed a separate task in which they received sweet, salty, sour, and neutral tastant solutions. We used both univariate and multivariate analysis techniques to compare the hemodynamic response to food pictures and direct taste stimulation.

## Results

### Behavioral Results

During the food pictures imaging scans, in which subjects saw pictures of a variety of sweet, sour, and salty foods as well as non-food objects (Figure 1; See Methods), subjects performed a picture repetition detection task with an average detection accuracy of 87.5%. Subjects also rated the pleasantness of those pictures, in a separate non-scanning task. Analysis of ratings from this food pleasantness rating task revealed a main effect of taste category (F = 11.1; p < 0.001). Sweet and Salty foods were rated as significantly more pleasant than Sour foods (p < 0.05), but no difference between sweet and salty foods (p = 0.59). During the non-scanning taste assessment task which followed, subjects rated the identity, pleasantness, and intensity of tasted delivered by our MR-compatible gustometer. There was a significant effect of tastant type on pleasantness ratings (F = 41.7; p < 0.001), with sweet rated as significantly more pleasant than salty or sour (sour: p < 0.02, salty: p < 0.001) and sour rated more pleasant than salty (p < 0.001), (see Table S1). There was also a main effect of tastant type on intensity ratings, with all tastants rated as more intense than neutral (p < 0.001). Subjects identified tastants during this task with an average accuracy of 97%.

**Figure 1:**
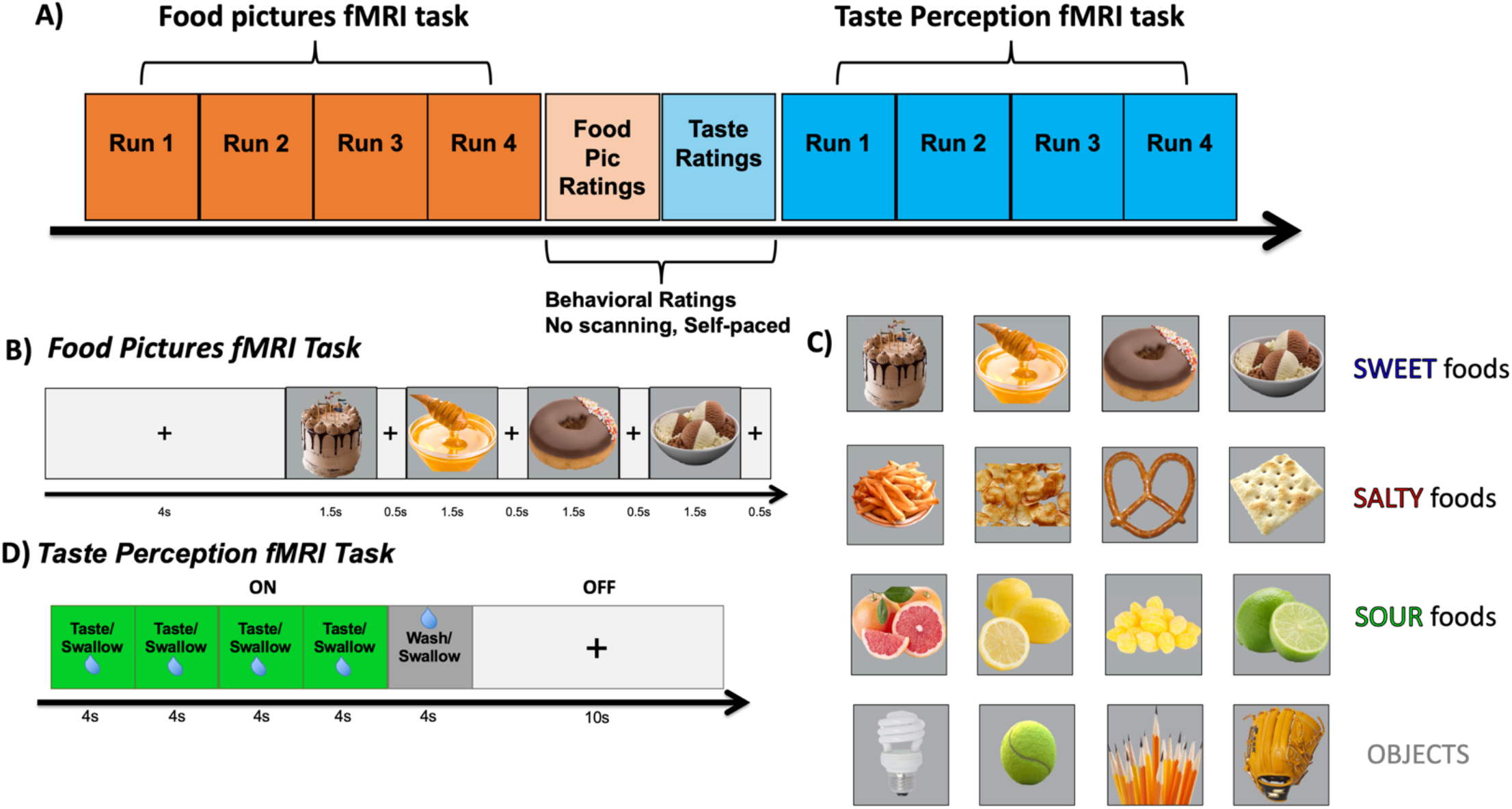
Experimental Design. A) An overview of the experimental session, during which participants performed Food Pictures and Taste Perception functional Magnetice Resonance Imaging (fMRI) task, separated by two non-scanning behavioral rating tasks. B & C) During the Food Pictures fMRI task, subjects viewed pictures of a variety of food and non-food objects within randomly ordered presentation blocks during scanning. Foods were categorized into predominantly Sweet, Sour, and Salty foods, as well as non-food familiar objects, selected on the basis of a series of prior experiments with a large online sample of participants. D) During the Taste Perception fMRI task, participants received sweet, sour, salty, and neutral tastant solutions, delivered in randomly ordered stimulus blocks, during scanning.

### Imaging Results – Univariate

#### Taste Perception task

Consistent with the previous results in an identical paradigm using tastant delivery during fMRI scanning (12), multiple clusters were identified within the insular cortex responsive to all tastants (sweet, sour, and salty) vs. the neutral solution. These clusters were located in bilaterally in the dorsal mid-insula, ventral anterior insula, and dorsal anterior insula (Figure 2A, Table 1). Importantly, the dorsal anterior insula cluster was located caudally to the most anterior areas of the insula, which have been shown to exhibit a more domain general role in task-oriented focal attention(19, 20). Beyond the insula, the bilateral regions of the ventral thalamus, postcentral gyrus, the cerebellum, and piriform cortex, and a region of the right putamen responded more to tastants relative to the neutral solution (Table 1).

**Figure 2:**
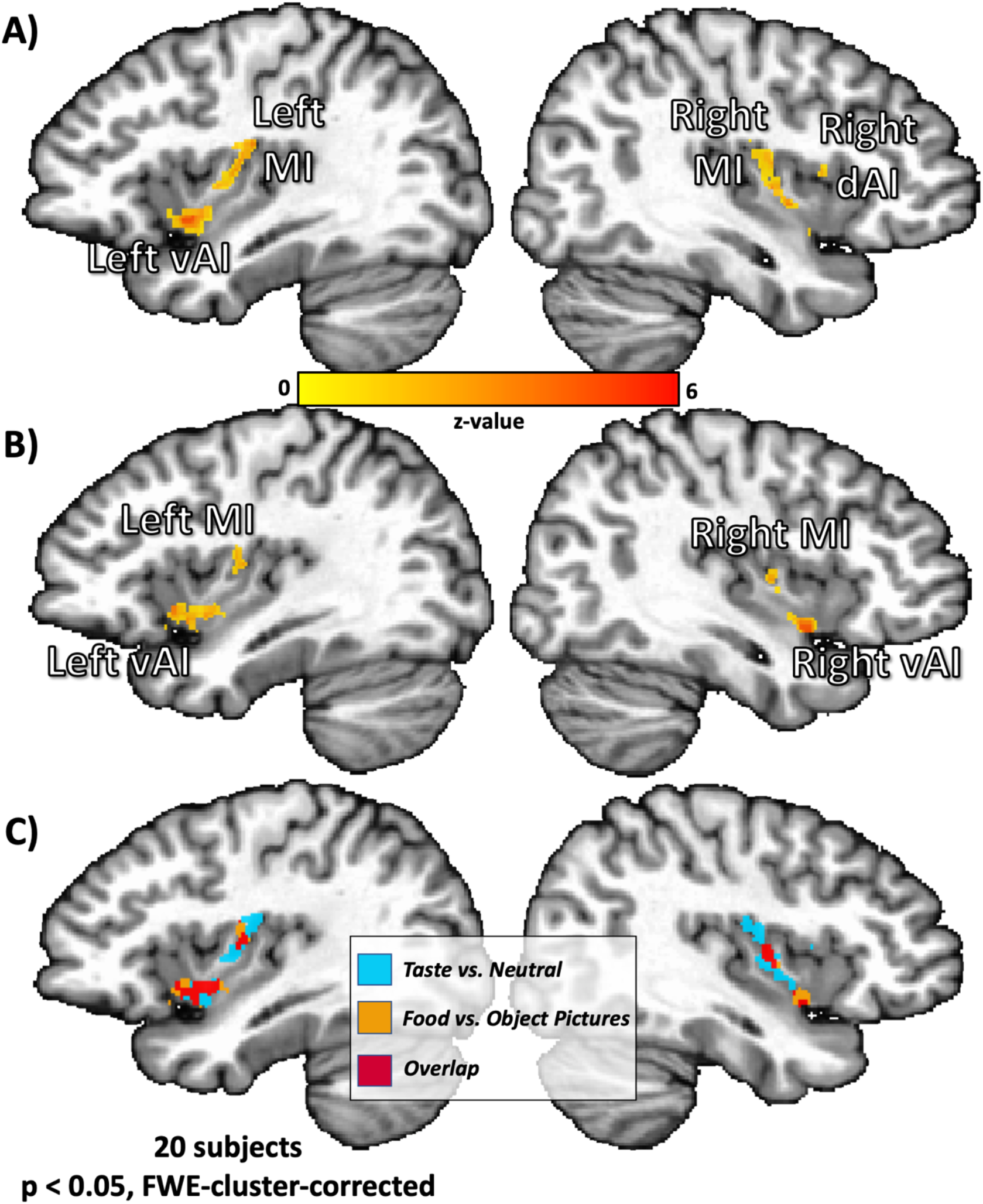
Bilateral regions of the ventral anterior and dorsal mid-insula are responsive to taste perception and viewing pictures of food. Univariate analysis results from both imaging tasks. A) All tastes (sweet, sour, and salty) vs. the neutral tastant activated bilateral regions of the dorsal mid insula (MI), as well as left ventral anterior (vAI) and right dorsal anterior insula (dAI). B) All food vs. object pictures activated bilateral mid-insula and ventral anterior insula, as well as left orbitofrontal cortex (not pictured). C) An overlap of the maps from (A) and (B) revealed bilateral ventral anterior and mid-insula regions co-activated by food pictures and taste. FWE - Family-wise Error

**Table 1.** Brain regions exhibiting significant a significant response to food vs. object pictures and taste vs. neutral tastes.

| Anatomical Location | Peak Coordinates |  |  | Peak Z | Cluster P | Volume, mm<sup>3</sup> |
| --- | --- | --- | --- | --- | --- | --- |
|  | X | Y | Z |  |  |  |
| Taste Perception task (All Tastes vs. Neutral) |  |  |  |  |  |  |
| R Cerebellum (VI) | -15 | 57 | -17 | 5.58 | <0.001 | 1,272 |
| L Dorsal Mid-Insula | 31.8 | 10 | 16.6 | 5.7 | <0.001 | 1,061 |
| R Thalamus | -11.4 | 17.4 | 2.2 | 5.93 | <0.001 | 949 |
| L Cerebellum (VI) | 9 | 54.6 | -15.8 | 5.72 | <0.001 | 8445 |
| R Mid-Insula | -37.8 | 6.6 | 4.6 | 5.32 | <0.001 | 835 |
| L Thalamus | 7.8 | 18.6 | -0.2 | 5.41 | <0.001 | 779 |
| R Postcentral Gyrus | -57 | 15 | 21.4 | 4.58 | <0.001 | 603 |
| L Ventral Anterior Insula | 36.6 | -7.8 | -5 | 4.48 | <0.001 | 353 |
| L Postcentral Gyrus | 55.8 | 10.2 | 17.8 | 4.6 | <0.01 | 252 |
| R Putamen | -24.6 | -1.8 | 8.2 | 4.37 | <0.02 | 133 |
| R Lingual Gyrus | -7.8 | 59.4 | 4.6 | 4.65 | <0.02 | 131 |
| R Precentral Gyrus | -59.4 | -0.6 | 17.8 | 4.48 | <0.02 | 123 |
| R Piriform Cortex / Amygdala | -24.6 | 6 | -11 | 4.49 | <0.03 | 98 |
| L Cuneus | 3 | 61.8 | 10.6 | 3.98 | <0.05 | 85 |
| R Dorsal Anterior Insula | -36.6 | -9 | 9.4 | 4.01 | <0.07 | 71 |
| Food Pictures task (All Food vs. Object Pictures) |  |  |  |  |  |  |
| Visual Cortex – Calcarine Gyrus | -5.4 | 81 | -7.4 | 5.2 | <0.001 | 3,166 |
| L Ventral Anterior Insula | 34.2 | -13.8 | -6.2 | 5.82 | <0.001 | 422 |
| L Area V3B | 23.4 | 78.6 | 14.2 | 4.6 | <0.001 | 204 |
| R Ventral Anterior Insula | -37.8 | -5.4 | -8.6 | 5.23 | <0.01 | 192 |
| L Orbitofrontal Cortex (BA11m) | 25.8 | -3.4 | -9.8 | 5 | <0.01 | 175 |
| L Dorsal Mid-Insula | 33 | 7.8 | 13 | 4.56 | <0.03 | 135 |
| R Area V3CD | -28.2 | 79.8 | 16.6 | 4.12 | <0.04 | 114 |
| R Mid-Insula | -37.8 | 5.4 | 7 | 6.65 | <0.10 | 60 |

#### Food Pictures task

A significant response for all food vs. object pictures was observed in bilateral regions of the mid-insula and ventral anterior insula, as well in the left lateral orbitofrontal cortex (area BA11m) (Figure 2B). These results are consistent with previous neuroimaging studies and meta-analyses of food picture presentation(3, 5, 7, 8). Relative to object pictures, viewing pictures of food also elicited activity in multiple areas of visual cortex (V1 and V3; see Table 1).

#### Conjunction Analysis

A conjunction of the corrected contrast maps generated for the Food Pictures and Taste Perception tasks and identified bilateral clusters within mid-insula and ventral anterior insula which exhibit overlapping activation for all tastes (vs. neutral) and all food pictures (vs. object pictures) (Figure 2C).

### Imaging Results - Multivariate

Previous studies have shown that multivariate pattern analyses can be used to distinguish between the distributed activity patterns by which distinct tastes are represented within the insular cortex and the wider brain (12, 17, 18). Our next analyses sought to answer these questions: A) Does MVPA allow us to reliably decode the taste category of food pictures within taste responsive regions of the brain? B) In which regions of the brain can we reliably decode both taste quality and food picture category? C) Within those overlapping regions, can we cross-classify food picture categories by training on experienced taste quality?

#### Insula ROI Analyses

Within the bilateral mid-insula clusters, MVPA revealed reliable classification between sweet, salty, and sour tastants (Left: Accuracy = 63%; p = 0.002; Right: Accuracy = 67%; p < 0.001; chance level = 50%; Fig. 3; Table S2) and between pictures of sweet, salty, and sour foods (Left: Accuracy = 71%; p < 0.001; Right: Accuracy = 65%; p = 0.002). Within the anterior insula clusters, we observed a task-specific dissociation between the ROIs, as the dorsal anterior insula clusters discriminated between tastes (Left: Accuracy = 64%; p < 0.001; Right: Accuracy = 65%; p< 0.001) but not food pictures (Left: Accuracy = 57%; corrected p = 0.09; Right: Accuracy = 56%; corrected p = 0.14), and the left ventral anterior insula cluster discriminated between food pictures (Accuracy = 65%; p < 0.001) but not tastes (Accuracy = 55%; corrected p = 0.25; Table S2). We tested this effect within our set of 6 ROIs using a permutation-test based ANOVA model. We used a model which included the laterality of the ROIs (LR) and whether they were located dorsally or ventrally (DV). We observed a significant Task*DV interaction (p = 0.025), and Task*LR interaction (p = 0.012), with no effect of task (p > 0.99), DV (p=0.11), LR (p > 0.99), or task*LR*DV interaction (p = 0.58).

**Figure 3:**
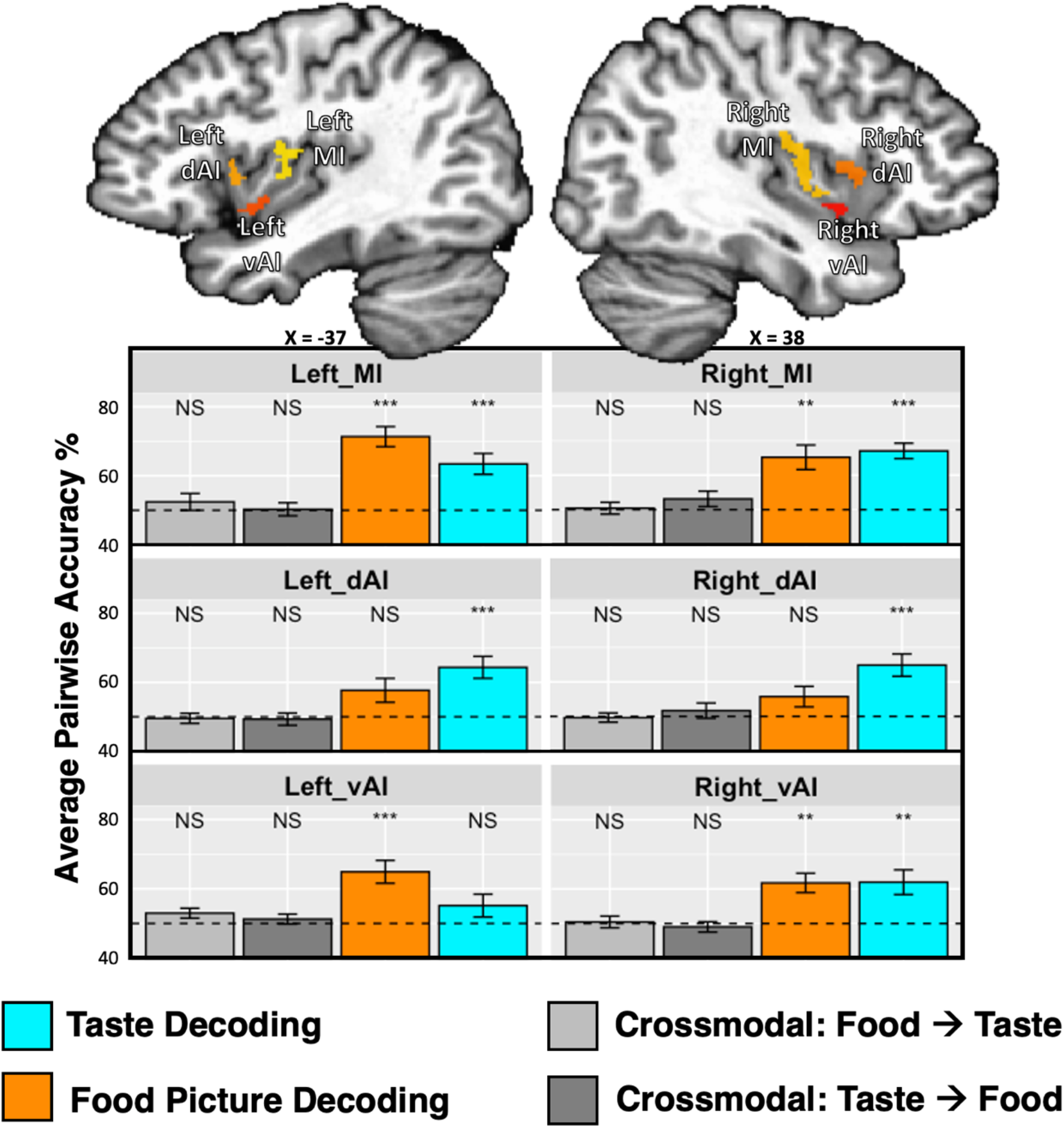
Multivariate pattern analyses reliably classify taste quality and food picture category within bilateral regions of the dorsal mid-insula. Taste-responsive regions of the bilateral insula, which were identified using the same taste paradigm within an independent dataset (Avery et al., 2020), were used as regions of interest for multivariate classification analyses performed on both imaging tasks. Within the bilateral dorsal mid-insula (MI), we were able to reliably classify both taste quality and food picture category, whereas we could classify only tastes within dorsal anterior insula (dAI). We could not reliably cross-classify food pictures by training on tastes, or vice versa. vAI – Ventral Anterior Insula.

#### Searchlight Analyses

The multivariate searchlight analysis using the Taste Perception data largely replicated the results of our previous study(12). Multiple, bilateral regions of the brain, including the mid-insula, dorsal anterior insula, somatosensory cortex, and piriform cortex, exhibited significant and above chance classification accuracy for discriminating between distinct tastes (see Table 2 for a comprehensive list of clusters).

**Table 2.** Brain regions where multivoxel patterns reliably discriminate between task categories.

| Anatomical Location | Peak Coordinates TLRC |  |  | Peak Z | Cluster P | Volume, mm³ |
| --- | --- | --- | --- | --- | --- | --- |
|  | X | Y | Z |  |  |  |
| Searchlight decoding: Taste Perception task |  |  |  |  |  |  |
| L inferior frontal / postcentral gyrus / insular cortex | 51 | -9 | 16.6 | 12.6 | <0.001 | 10,064 |
| R anterior and mid insula / postcentral Gyrus | -33 | -9 | 9.4 | 13.6 | <0.001 | 9,215 |
| R cerebellum / pons / parahippocampal gyrus / L fusiform gyrus | -7.8 | 41.4 | -20.6 | 9.1 | <0.001 | 6,304 |
| R medial thalamus | -9 | 16.2 | -1.4 | 9.5 | <0.02 | 1,472 |
| R hippocampus / parahippocampal gyrus / amygdala / cerebellum | -29.4 | 24.6 | -11 | 10.5 | <0.03 | 1,346 |
| R lateral orbitofrontal cortex (BA47) | -42.6 | -41.4 | -8.6 | 11.9 | <0.03 | 1,120 |
| R superior temporal sulcus | -43.8 | 6.6 | -15.8 | 9.5 | <0.03 | 921 |
| L middle frontal gyrus | 35.4 | -43.8 | 15.4 | 9.9 | <0.03 | 871 |
| L middle temporal gyrus | 42.6 | 6.6 | -20.6 | 9.4 | <0.03 | 840 |
| R piriform cortex | -40.2 | -1.8 | -21.8 | 9.6 | <0.03 | 835 |
| R middle frontal gyrus | -43.8 | -37.8 | 15.4 | 10.8 | <0.05 | 634 |
| R cerebellum | -17.4 | 55.8 | -18.2 | 9.5 | <0.05 | 591 |
| Searchlight decoding: Food Pictures task |  |  |  |  |  |  |
| Bilateral visual / ventral temporal cortex | 24.6 | 52.2 | -12.2 | 27.2 | <0.001 | 128,259 |
| L mid-insula | 37.8 | 9 | 11.8 | 12.5 | <0.001 | 2,053 |
| L post-central gyrus | 55.8 | 25.8 | 27.4 | 13.2 | <0.001 | 1,954 |
| L orbitofrontal cortex (ba11m) | 24.6 | -30.6 | -8.6 | 15.2 | <0.01 | 1,555 |
| R orbitofrontal cortex (ba11m) | -25.8 | -31.8 | -9.8 | 14.8 | <0.01 | 1,453 |
| R mid-Insula / ventral anterior insula | -37.8 | 5.4 | 7 | 11.9 | <0.01 | 1,203 |
| R post-central gyrus | -57 | 18.6 | 22.6 | 12.4 | <0.01 | 1,092 |
| R inferior frontal gyrus | -42.6 | -28.2 | 16.6 | 11.9 | <0.01 | 869 |
| L inferior frontal gyrus | 41.4 | -27 | 19 | 12.3 | <0.02 | 674 |
| L ventral anterior insula | 36.6 | -7.8 | -2.6 | 9.8 | <0.04 | 411 |
| L amygdala | 19.8 | 4.2 | -13.4 | 9 | <0.05 | 335 |
| Conjunction Food Pictures and Taste Decoding |  |  |  |  |  |  |
| L fusiform gyrus / parahippocampal gyrus | -37 | -46 | -24 | 3 | - | 1,448 |
| R parahippocampal gyrus | 21 | -31 | -18 | 3 | - | 817 |
| L postcentral gyrus | -57 | -15 | 14 | 3 | - | 508 |
| L dorsal mid-insula | -36 | -3 | 3 | 3 | - | 499 |
| R dorsal mid-insula | 40 | -7 | -0 | 3 | - | 219 |
| R fusiform gyrus | 28 | -52 | -22 | 3 | - | 195 |
| R fusiform gyrus | 34 | -40 | -24 | 3 | - | 161 |
| R postcentral gyrus | 58 | -14 | 20 | 3 | - | 133 |

The searchlight analysis performed on the Food Pictures task also identified multiple regions, including bilateral regions of the dorsal mid-insula, ventral anterior insula, postcentral gyrus (approximately located in the oral somatosensory strip), orbitofrontal cortex, and the left amygdala. Significant classification accuracy was also observed bilaterally in the inferior frontal gyrus and widespread regions of occipital cortex, stretching into both dorsal and ventral processing streams, including the parahippocampal gyrus, bilaterally (Figure 4, Table 2).

**Figure 4:**
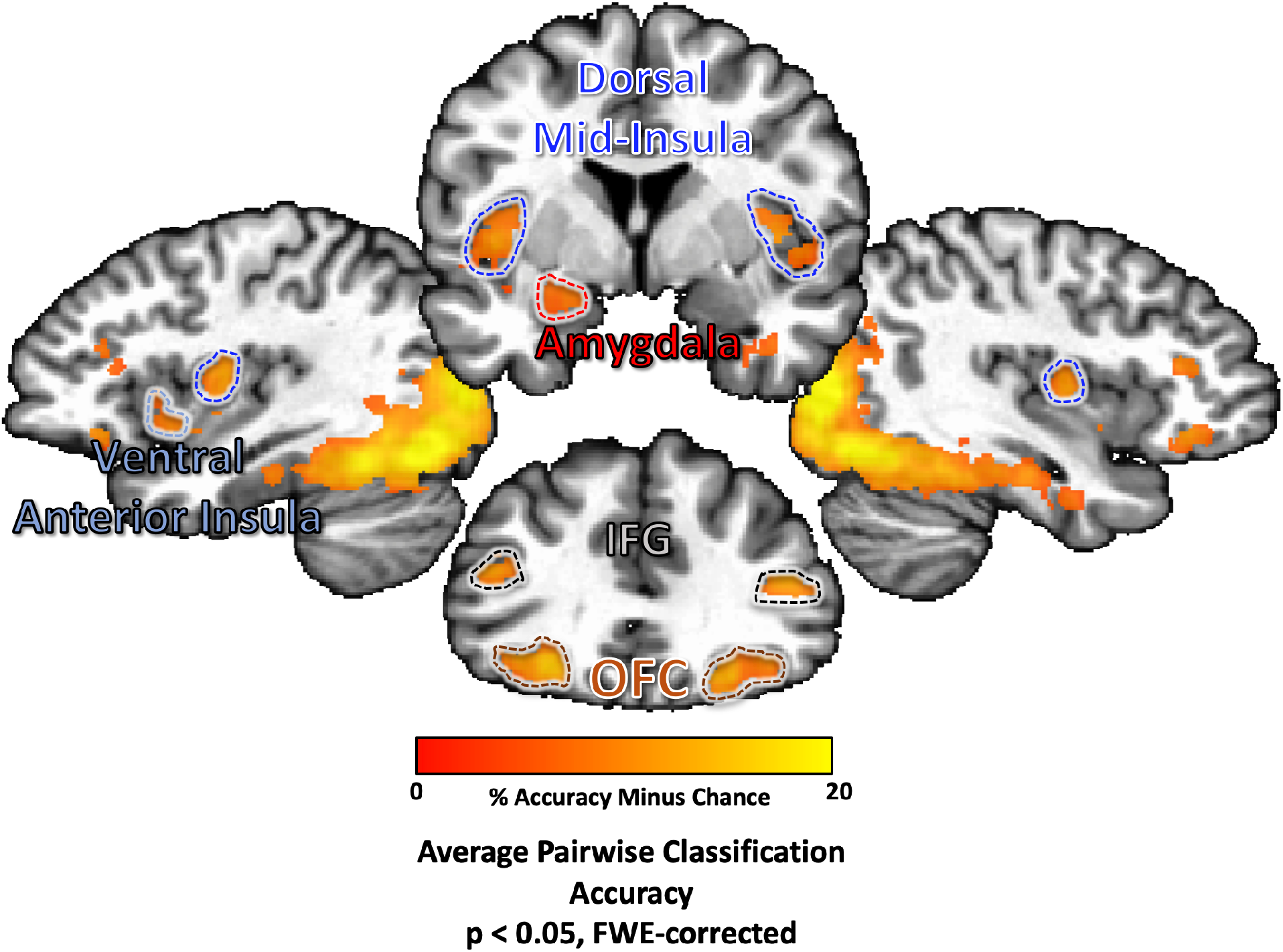
Multivariate pattern analyses classify food pictures according to taste category within brain regions involved in taste perception, arousal, and reward. Several regions of the brain were identified using a multivariate searchlight analysis trained to discriminate between pictures of sweet, salty, and sour foods. This included regions involved in processing the sensory and affective components of food, including bilateral regions of the dorsal mid-insula and ventral anterior insula, as well as the bilateral postcentral gyrus, the bilateral orbitofrontal cortex (OFC), inferior frontal gyrus (IFG), and the left amygdala.

A conjunction of the searchlight classification accuracy maps for both tasks identified bilateral regions of dorsal mid-insular cortex, the post-central gyrus, the fusiform gyrus, and parahippocampal gyrus (Figure 5, Table 2).

**Figure 5:**
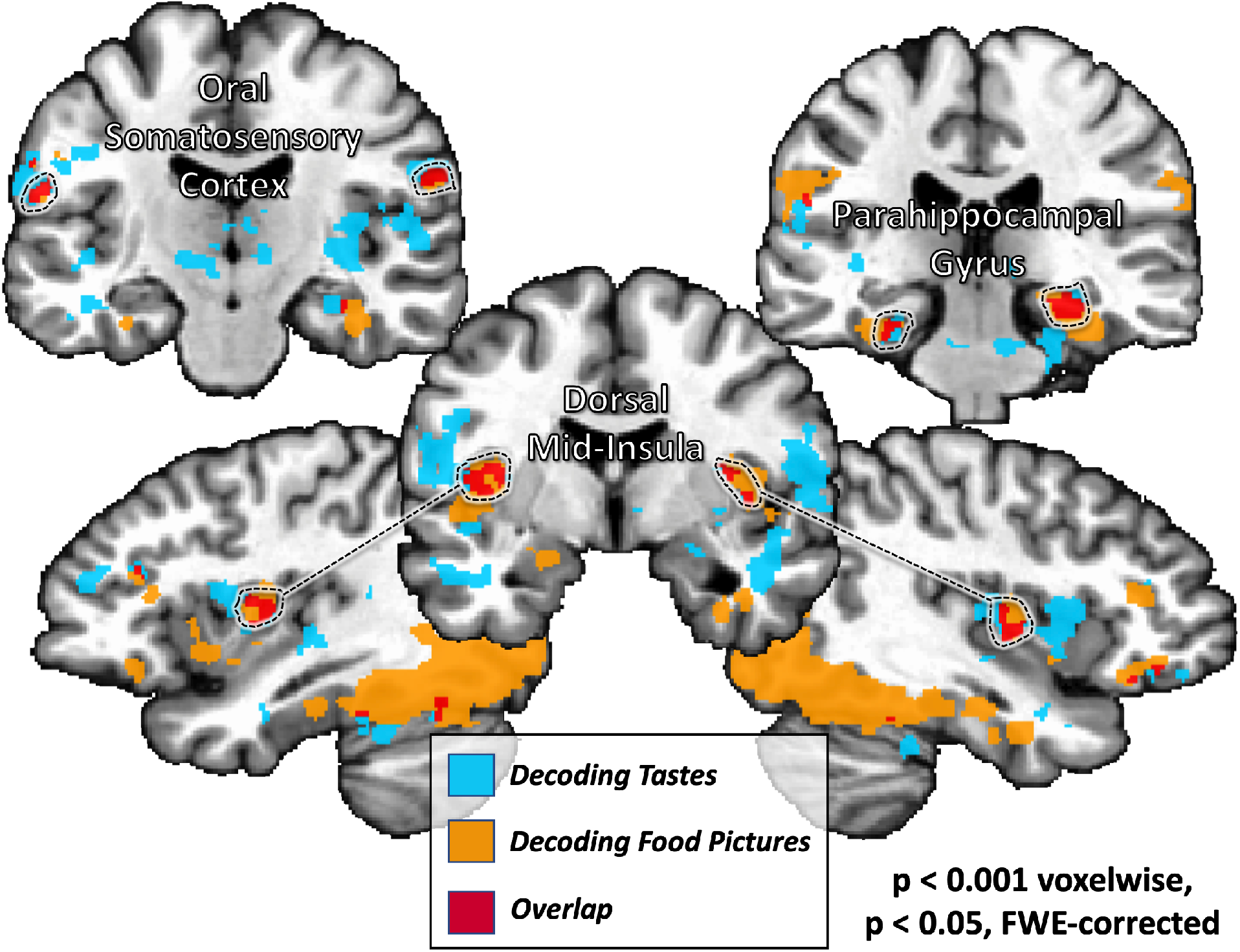
A common set of brain regions supporting information about taste quality and food picture category. A conjunction of multivariate searchlight maps performed on both Taste Perception and Food Pictures tasks identifies a set of regions which reliably discriminate between both taste quality and food picture category. This set of regions included sensory cortical areas within the mid-insular cortex and somatosensory cortex, as well as regions of the ventral occipitotemporal object processing stream.

#### Object Pictures Analyses

Using a whole-brain multivariate searchlight analysis, we were able to reliably classify the non-food object pictures within widespread areas of the occipitotemporal cortex, including primary visual cortex and much of the ventral visual processing stream (Figure S2a). Critically, the effects were limited to these visual processing regions of the brain and did not include any of the other areas involved in classifying food pictures, such as the insula, OFC, or amygdala. To confirm these results, we also ran an ROI analysis within our 6 insula ROIs and determined that the object picture classification accuracy was no greater than chance (Figure S2b).

#### Crossmodal Decoding

An SVM decoder was used to classify food picture categories using taste categories from the Taste Perception task as training data, and vice-versa. This analysis failed to identify any evidence of cross-classification within the insula ROIs, though the Right Mid-Insula ROI did exhibit a non-significant trend (Accuracy: 56%, FDR-corrected p = 0.11) when training on taste and testing on food pictures (Figure 3, Table S2). A follow-up permutation-test based ANOVA within the insula ROIs identified a significant effect of modality (p < 0.001), which indicates that decoding models trained and tested *within*-modality were significantly more accurate than decoding models trained and tested *across* modality.

Following this, we examined the similarity of multivariate patterns produced by food pictures and tastes in the mid-insula, both within modality and between modality. Using splithalf correlations of food picture and taste data, we determined that patterns produced by both of these tasks had high within-modal similarity (Figure S3). However, the patterns produced by these different tasks were highly dissimilar from each other, as the between-modality similarity was no different from zero, as well as significantly less than within modal similarity (ANOVA modality effect: p < 0.001; Figure S3).

Furthermore, we also examined the distribution of voxel weights generated by our SVM models to separately classify tastes and food pictures within our mid-insula ROIs. These SVM weights were generated following a backward-to-forward model transformation procedure, which allows these parameters to be more readily interpretable in terms of the brain processes under study (21). These parameters then can be used to indicate the voxels within each ROI which are most informative for predicting a particular taste or food picture category. We calculated the spatial correlation of these voxel weights on a subject-by-subject basis, both within modality (split-half) and between modality, and examined the average correlations at the group level. Again, we observed significant within-modality correlations, but the between modal correlations were again no different than zero, as well as significantly less than within-modal correlations (ANOVA modality effect: p < 0.001; Figure S4). This indicates that the multivariate decoding models relied on independent sets of voxels when classifying food pictures and tastes.

We also performed cross-modal classification analyses within a whole-volume searchlight and failed to observe any regions exhibiting significant cross-classification accuracy, both when training on tastes and training on pictures.

### Pleasantness analyses

To identify the effect of self-reported pleasantness ratings on the hemodynamic response to tastants and pictures, a whole-volume t-test of parametrically modulated hemodynamic response functions, as well as a whole-volume linear mixed-effects regression model, were employed. After correction for multiple comparisons, neither approach was able to identify any brain regions exhibiting a reliable relationship between pleasantness ratings and tastant response, for either task. At the ROI level, we examined the effect of pleasantness ratings on the response to food pictures within the taste-responsive clusters of the insula. We identified an effect of taste (F = 3.3; p = 0.04) and an effect of region (F = 5.1; p = 0.002), but no effect of pleasantness ratings (p = 0.18).

## Discussion

The objective of this study was to determine whether taste responsive regions of the brain represent the inferred taste quality of visually presented foods. The subjects within this study viewed a variety of food pictures which varied categorically according to their dominant taste: sweet, sour, or salty. While viewing these pictures, the subjects performed a picture-repetition detection task, a task which was orthogonal to the condition of interest in this study. In a separate task, those same subjects also directly experienced sweet, sour, and salty tastes, delivered during scanning. Viewing food pictures and directly experiencing tastes activated overlapping regions of the dorsal mid-insula and ventral anterior insula. This finding is in agreement with neuroimaging meta-analyses of taste and of food picture representation, which have implicated both regions in these separate functional domains(7, 8, 22, 23).

### Multivariate patterns representing tastes and food pictures

Based upon previous evidence that the insula represents taste quality by distributed patterns of activation within taste-responsive brain regions, rather than topographically(12, 17, 18), we sought to identify whether this region uses a similar distributed activation pattern to represent the taste quality of visually presented foods. Using multivariate pattern analysis, we were able to build upon those previous findings by demonstrating that the taste categories associated with food pictures could be reliably discriminated in taste-responsive regions of the ventral anterior and dorsal mid-insula. Moreover, within the bilateral dorsal mid-insula specifically, we were able to reliably classify both the taste quality of tastants and the taste category of food pictures. These results suggest that viewing pictures of food does indeed trigger an automatic retrieval of taste property information within taste-responsive regions of the brain and most importantly, this retrieved information is detailed enough to represent the specific taste qualities associated with visually presented foods.

These results demonstrate how higher-order inferences derived from stimuli in one modality (in this case vision) can be represented in brain regions typically thought to represent only low-level information about a different modality (in this case taste). Broadly, these results echo previous neuroimaging findings on multisensory processing within vision and audition, which demonstrate that early sensory cortical areas are able to represent the inferred sensory properties of stimuli presented via another sensory channel(24, 25). Taken together, these findings are consistent with claims that both higher-order and presumptively unimodal areas of neocortex are fundamentally multisensory in nature (26).

Using a multivariate searchlight approach, we were also able to identify a broad network of regions, outside of the insular cortex, within which we were able to reliably classify the taste quality of visually presented foods. Those regions included the left amygdala and bilateral orbitofrontal cortices, regions previously observed in food picture neuroimaging studies(5, 7, 8) and typically associated with food-related affect and reward (27–29). We also observed significant classification accuracy for food picture categories within the bilateral postcentral gyrus, inferior frontal gyrus, and widespread regions of the visual cortex. A control searchlight analysis, run on the pictures of non-food objects from this task, was able to classify the object pictures within these same areas of visual cortex, suggesting that these regions were carrying information about low-level visual features of the picture stimuli. Critically though, the effects did not include any of the other areas involved in classifying food pictures, such as the insula, orbitofrontal cortex, or amygdala. Only when decoding pictures of food could we reliably classify within brain regions involved in the experience of taste.

Within the human object recognition pathway, lower level visual cortex regions pass on information to ventral temporal areas associated with object recognition which then send that information forward to the OFC, ventral striatum, and amygdala(30, 31). Importantly, the amygdala and OFC also sit directly downstream of the insula in the taste pathway(31, 32) and play a role in appetitive and aversive behavioral responses to taste, such as conditioned taste aversion(33). The OFC has been demonstrated to represent the reward value of food cues(27–29). The OFC, in concert with the amygdala and mediodorsal thalamus, is thought to represent the dynamic value of environment stimuli and sensory experiences, informed by the body’s current state(34). Recent rodent studies have shown that the amygdala directly signals gustatory regions of the insula in response to food predictive cues, in a manner which is specifically gated by hypothalamic signaling pathways and is thus differentially responsive to states like hunger or thirst(35). Thus, the amygdala sits in a position to serve as a neural relay linking the taste system with the object recognition system, which allows us to infer the homeostatically relevant properties of visually perceived food stimuli within our environment.

We ran a comparable searchlight analysis of data from the taste task and performed a conjunction of the classification maps from the two tasks. We observed that, beyond the dorsal mid-insula, several regions of this network also reliably discriminated between both food picture taste categories and tastes. This set of regions included sensory cortical areas such as the bilateral dorsal mid-insula and bilateral regions of post-central gyrus at the approximate location of oral somatosensory cortex. This set also included bilateral regions of the parahippocampal gyrus and fusiform gyrus, regions of the ventral visual stream which represent high-level object properties. Interestingly, previous neuroimaging studies have also identified odor-evoked effects in higher order visual regions, such as the fusiform gyrus (36, 37), which suggest that olfactory regions directly exchange information about stimulus identity with this region of extended visual cortex. Another recent study also identified a specific region of the fusiform gyrus associated with accuracy at estimating the energy density of visually depicted foods (38). The observation that this set of regions support category-specific patterns of activity for both tastes and food pictures lends further support to the idea that the fusiform gyrus plays a role in representing higher-order information about food, which may be used to guide value-based decision making (38).

### Crossmodal Decoding

We also examined the possibility that the food-picture-evoked activity patterns within taste-responsive insula regions were similar to the taste-evoked patterns, by using a cross-modal classification analysis. Our results suggest that this is not the case, as we were unable to reliably discriminate between food picture categories after training using the corresponding taste categories during the taste task. Further analysis of these results indicated that not only were the multivariate patterns produced by tastes and pictures completely unrelated, but the decoding models relied most heavily on independent sets of voxels when classifying food pictures or tastes. These results are partly in keeping with previous studies which have generated similar null results when attempting cross-modal classification within primary sensory cortices(24, 25).

There are several possible explanations for our failure to cross-decode inferred and experienced tastes. One possibility is that inferred and experienced tastes activate different neural populations within the mid-insula, even when the inferred and experienced tastes represent the same basic information. In that view, inferring a taste (e.g., salty) and experiencing it may both activate the same region but with a different response pattern. This could provide a vital mechanism by which inferred and experienced sensations are kept distinct at the neural level, in order to prevent inappropriate physiological or behavioral responses to inferred sensations.

A related possibility is that the neural signals conveying taste and picture information are relayed to gustatory cortex via different neural pathways, and thus may terminate in different cortical layers of the insula. Currently, even ultra-high-resolution fMRI imaging, such as was employed for this study, would be unable to distinguish distinct populations of intermingled neurons at sub-voxel resolution. However, future neuroimaging studies employing more indirect imaging paradigms such as fMRI-Adaptation (39, 40), in combination with emerging techniques for laminar-level fMRI-imaging (41, 42), might be used to discriminate these sub-voxel level responses. Along these lines, if the inferred tastes of food pictures can be shown to selectively adapt the response to directly experienced tastes, this would indicate the activation of a shared population of neurons activated by both modalities.

Another possibility is that the dissimilarity of these patterns reflects the relative experiential distance between actual consumption of a taste and merely viewing a stimulus that is predictive of consumption. Due to the limitations of picture stimulus presentation during scanning, we were unable to replicate many of the salient features of real-world foods which are present during our everyday interactions, such as their relative size and graspability. Previous studies have demonstrated that the physical presence of a food, compared to viewing a picture of it, increases subjects’ willingness to pay for that food and the expected satiety upon its consumption (43, 44). Additionally, cephalic phase responses such as insulin release and salivation all greatly increase with our degree of sensory exposure to a food, going from sight and smell all the way to initial digestion (45). These studies suggest that the format in which a food is presented affects both our valuation of it as well as our conceptual representation of its sensory properties. Potentially, this greater degree of sensory exposure to foods would be reflected in greater multivariate pattern similarity within gustatory cortical regions.

Alternatively, the inferences generated by the depicted foods may represent a more complex variety of properties than simply taste, including overall flavor (i.e. the combination of taste and smell), texture, and fat content/appetitiveness. Indeed, previous neuroimaging studies have demonstrated the involvement of the mid and ventral anterior insula in representing flavor, fat content, and the viscosity of orally delivered solutions (46–48). Future studies, applying such techniques as representational similarity analysis, along with an appropriately designed stimulus set, could identify to what degree these various food-related properties contribute to the multivariate patterns present within the taste-responsive regions of the insula.

### Functional specialization within the insula

Interestingly, through our multivariate analyses, we observed a task-related functional dissociation within the anterior insular cortex, as the dorsal anterior insula discriminated between basic tastes but not food pictures. This suggests that taste-related information might be relayed within the insula along separate dorsal and ventral routes from the dorsal mid-insula to the respective regions of anterior insula, which in turn transmit this information to their associated functional networks. Meta-analyses of multiple neuroimaging studies have indicated that the dorsal and ventral regions of the anterior insula are associated with distinctly different domains of cognition(20, 49). The dorsal anterior insula has high connectivity with frontoparietal regions and is associated with cognitive and goal-directed attentional processing(19, 20, 49). In contrast, the ventral anterior insula has a high degree of connectivity with limbic and default-mode network regions and is much more implicated in social and emotional processing(20, 49). Ventral anterior insula is also thought to serve as a link between the gustatory and olfactory systems, whose key function would be to integrate taste and smell to produce flavor(13, 46). The results of the present study also strongly link the ventral anterior insula with the ventral occipitotemporal pathway involved in object recognition. This dorsal/ventral functional division of the anterior insula would thus potentially mirror the action/stimulus division of the dorsal and ventral striatum(50). Relatedly, we also observed some effect of laterality within the insula, with slightly greater accuracy for taste decoding in the right than the left insula. This laterality finding seems to concur with prior evidence of a specialization of the right insula for processing taste concentration (51).

### Food and taste cues as interoceptive predictions

In addition to its role in gustatory perception, the insula also serves as primary interoceptive cortex, receiving primary visceral afferents from peripheral vagal and spinothalamic neural pathways(52). The involvement of the insula, the dorsal mid-insula in particular, in interoceptive awareness has been well established in human neuroimaging studies(9, 53). Indeed, previous neuroimaging studies have provided evidence that the dorsal mid-insula serves as convergence zone for gustatory and interoceptive processing(9, 10), evidence strongly supported by studies of homologous regions of rodent insular cortex(54, 55). Indeed, the activity of dorsal mid-insula to food images seems acutely sensitive to internal homeostatic signals of energy availability(4, 56). According to recent optical imaging studies in rodents, food predictive cues transiently modify the spontaneous firing rates of insular neurons, pushing their activity toward a predicted state of satiety(57). Thus the automatic retrieval of taste property information for food pictures can be understood within an interoceptive predictive-coding framework(58), in which viewing pictures of homeostatically-relevant stimuli modifies the population-level activity of gustatory/interoceptive regions of the insula, in a manner specific to that food’s predicted impact upon the body.

### The role of Pleasantness

We also attempted to minimize and account for the role of pleasantness within the present study, as a way of focusing specifically on taste quality within both tasks, as opposed to merely the perceived pleasantness of tastants of food pictures. To this end, we used mild concentrations of our sweet, sour, and salty tastants, as we did in our previous study(12). We also used sets of food pictures which had all been rated as highly pleasant by an online sample (See Supplemental Materials). Nevertheless, subjects reported that tastes and food pictures did differ in pleasantness. We examined the effect of pleasantness upon responses to tastes and food pictures using separate approaches, one in which participants’ pleasantness ratings were used to account for trial-by-trial variance (i.e., amplitude modulation regression) and one in which ratings were used to account for any remaining variance at the group level. Consistent with our previous study(12), neither approach showed an effect of pleasantness on the hemodynamic response to tastes or food pictures, at the whole-volume or at the ROI level. Importantly, within those insula ROIs, we were able to reliably discriminate between pictures of salty and sweet foods (see Figure S2), which did not differ in pleasantness, thus suggesting that pleasantness did not account for the observed results in this study.

## Conclusion

The goal of this study was to determine whether taste-responsive regions of the insula also represent the specific inferred taste qualities of visually presented foods and whether they do so using reliably similar patterns of activation. We were able to reliably classify the taste category of food pictures within multiple regions of the brain involved in taste perception and food reward. We additionally identified several regions of the brain, including the bilateral dorsal mid-insular cortex, in which we were able to decode both food picture category and the taste quality of tastants delivered during another task. However, we were unable to reliably cross-decode food picture category by training on the corresponding taste quality within any of these regions. This suggests that while these regions are able to represent both inferred and experienced taste quality, they do so using distinct patterns of activation.

## Materials and Methods

### Participants

We recruited 20 healthy subjects (12 female) between the ages of 21 and 45 (Average(SD): 26(7) years). Ethics approval for this study was granted by the NIH Combined Neuroscience Institutional Review Board under protocol number 93-M-0170. The institutional review board of the National Institutes of Health approved all procedures, and written informed consent was obtained for all subjects. Participants were excluded from taking part in the study if they had any history of neurological injury, known genetic or medical disorders that may impact the results of neuroimaging, prenatal drug exposure, severely premature birth or birth trauma, current usage of psychotropic medications, or any exclusion criteria for magnetic resonance imaging (MRI).

### Experimental Design

All fMRI scanning and behavioral data was collected at the NIH Clinical Center in Bethesda, MD. Participant scanning sessions began with a high-resolution anatomical reference scan followed by an fMRI scan, during which subjects performed our Food Pictures task (Figure 1). This scan was followed by a short non-scanning task in which participants rated the pleasantness of several of the food pictures they had viewed during the previous task. Next, participants performed a non-scanning Taste Assessment task, during which they rated the tastants on the pleasantness, identity, and intensity. Finally, participants performed our Taste Perception task during fMRI scanning. The methods used for the Taste Perception task and taste assessment were nearly identical to those used in our previous study(12). Importantly, all tasks requiring tastant delivery were performed after food picture tasks, to avoid the possibility of any carryover or priming effects of the tastants upon the response to the food pictures.

### Food Pictures fMRI task

During this task, participants viewed images of various foods and non-food objects. Pictures were presented sequentially, with four pictures shown per presentation block. Each block consisted of four pictures of either sweet, sour, or salty foods or of specific types of non-food familiar objects. Pictures were presented at the center of the display screen against a gray background. Within a presentation block, pictures were presented for 1500ms, followed by a 500ms interstimulus interval (ISI), during which a fixation cross ‘+’ appeared on the screen. Another four-second ISI followed each presentation block (see Figure 1). Presentation blocks were presented in a pseudo-random order by picture category, with no picture category presented twice in a row.

The food types presented during this task were twelve foods selected and rated to be predominantly sweet (cake, honey, donuts, ice cream), sour (grapefruit, lemons, lemon candy, limes), or salty (chips, fries, pretzels, crackers), by groups of online participants recruited through Amazon Mechanical Turk (for details, see Supplemental Materials: Online Experiments section, also see Figure S1). The selection criteria for these foods was that they be clearly recognizable, pleasant, and strongly characteristic of their respective taste quality (see Figure S1). Non-food objects were familiar objects - basketballs, tennis balls, lightbulbs (fluorescent and incandescent), baseball gloves, flotation tubes, pencils, and marbles - which roughly matched the shape and color of the pictured foods. In total, participants of the fMRI study viewed 28 unique exemplars of each type of the 12 foods (336 total) and 14 unique exemplars of the non-food objects (112 total).

During half of the presentation blocks, one of the food or object pictures was repeated, and participants were instructed to press a button on a handheld fiber optic response box whenever they identified a repeated picture. Blocks with repetition events were evenly distributed across picture categories (sweet, salty, sour, and object pictures), such that half the blocks of each category contained a repetition. Repetition blocks were also evenly distributed across food and object types, such that each food type was used in a repetition event four times and each object type was used twice. Eight presentation blocks for each picture category (sweet, salty, sour, & objects) were presented during each run of the imaging task (32 total). Each of the four imaging runs lasted for 388 seconds (6 minutes, 28 seconds). For MVPA analysis, each run was split into two run segments, which allowed us to use a total of eight run segments for subsequent MVPA analyses.

### Food pleasantness rating task

Following the fMRI Food Pictures task, participants were then asked to perform a separate task in which they rated the pleasantness of the food pictures they had seen during the imaging task. Three exemplars of each type of food picture (36 total) were presented in random order against a gray background. Participants were asked to indicate on a 0 (not pleasant at all) to 10 (extremely pleasant) scale, using the handheld response box, how pleasant it would be to eat the depicted food at the present moment. These rating periods were self-paced, and no imaging data were collected during this task.

### Taste assessment

Participants next completed a taste assessment task, during which they received 0.5mL of a tastant solution delivered directly onto their tongue by an MR-compatible tastant delivery device. Four types of tastant solutions were delivered during the taste assessment: Sweet (0.6M sucrose), Sour (0.01M citric acid), Salty (0.20M NaCl), and Neutral (2.5mM NaHCO3 + 25mM KCl). Subjects were then prompted to use the handheld response to indicate the identity of the tastant they received, as well as the pleasantness and intensity of that tastant. Following these self-paced rating periods, the word “wash” appeared on the screen and subjects received 1.0mL of the neutral tastant, to rinse out the preceding taste. This was followed by another (2s) prompt to swallow. A four-second fixation period separated successive blocks of the taste assessment task. Tastants were presented five times each (20 blocks total), in random order. Altogether, this session lasted between five to seven minutes. All tastants were prepared using sterile lab techniques and USP-grade ingredients by the NIH Clinical Center Pharmacy.

### Gustometer Description

A custom-built pneumatically-driven MRI-compatible system delivered tastants during fMRI-scanning(4, 9–12). Tastant solutions were kept at room temperature in pressurized syringes and fluid delivery was controlled by pneumatically-driven pinch-valves that released the solutions into polyurethane tubing that ran to a plastic gustatory manifold attached to the head coil. The tip of the polyethylene mouthpiece was small enough to be comfortably positioned between the subject’s teeth. This insured that the tastants were always delivered similarly into the mouth. The pinch valves that released the fluids into the manifold were open and closed by pneumatic valves located in the scan room, which were themselves connected to a stimulus delivery computer that controlled the precise timing and quantity of tastants dispensed to the subject during the scan. Visual stimuli for behavioral and fMRI tasks were projected onto a screen located inside the scanner bore and viewed through a mirror system mounted on the head-coil. Both visual stimulus presentation and tastant delivery were controlled and synchronized via a custom-built program developed in the PsychoPy2 environment.

### Taste Perception Task

During the Taste Perception fMRI task, the word ‘Taste’ appeared on the screen for 2s, and subjects received 0.5mL of either a sweet, sour, salty, or neutral tastant. Next, the word “swallow” appeared on the screen for 2s, prompting subjects to swallow. These taste and swallow periods occurred four times in a row, with the identical tastant delivered each time. Following these four periods, the word “wash” appeared on the screen and subjects received 1.0mL of the neutral tastant, to rinse out the preceding tastes. This was followed by another (2s) prompt to swallow. In total, these taste delivery blocks lasted 20 seconds. These taste delivery blocks were followed by a 10-second ISI, during which a fixation cross ‘+’ was presented on the center of the screen. Four sweet, salty, sour, and neutral taste delivery blocks (16 total) were presented in random order throughout each run of this task. Each run contained a 4-second initial fixation period and another 6-second fixation period halfway through the run, for a total of 490 seconds/run (8 minutes, 10 seconds). Participants completed 4 runs of the Taste Perception task during one scan session. As with the Food Pictures task, each run of this task was split into two run segments. This allowed us to use a total of eight run segments for subsequent MVPA analyses.

### Imaging Methods

fMRI data was collected at the NIMH fMRI core facility at the NIH Clinical Center using a Siemens 7T-830/AS Magnetom scanner and a 32-channel head coil. Each echo-planar imaging (EPI) volume consisted of 58 1.2-mm axial slices (echo time (TE) = 23 ms, repetition time (TR) = 2000 ms, flip angle = 56 degrees, voxel size = 1.2 × 1.2 × 1.2 mm^3^). A Multi-Band factor of 2 was used to acquire data from multiple slices simultaneously. A GRAPPA factor of 2 was used for inplane slice acceleration along with a 6/8 partial Fourier k-space sampling. Each slice was oriented in the axial plane, with an Anterior-to-Posterior phase encoding direction. Prior to task scans, a 1-minute EPI scan was acquired with the opposite phase encoding direction (Posterior-to-Anterior), which was used for correction of spatial distortion artifacts during preprocessing (see Image Preprocessing). An ultra-high resolution MP2RAGE sequence was used to provide an anatomical reference for the fMRI analysis (TE = 3.02 ms, TR = 6000 ms, flip angle = 5 degrees, voxel size = 0.70 × 0.70 × 0.70 mm).

### Image Preprocessing

All fMRI pre-processing was performed in AFNI (http://afni.nimh.nih.gov/afni). The FreeSurfer software package (http://surfer.nmr.mgh.harvard.edu/) was additionally used for skullstripping the anatomical scans. A de-spiking interpolation algorithm (AFNI’s 3dDespike) was used to remove transient signal spikes from the EPI data, and a slice timing correction was then applied to the volumes of each EPI scan. The EPI scan acquired in the opposite (P-A) phase encoding direction was used to calculate a non-linear transformation matrix, which was used to correct for spatial distortion artifacts. All EPI volumes were registered to the very first EPI volume of the Food Pictures task using a 6-parameter (3 translations, 3 rotations) motion correction algorithm, and the motion estimates were saved for use as regressors in the subsequent statistical analyses. Volume registration and spatial distortion correction were implemented in the same non-linear transformation step. A 2.4mm (2-voxel width) FWHM Gaussian smoothing kernel was then applied to the volume-registered EPI data. Finally, the signal intensity for each EPI volume was normalized to reflect percent signal change from each voxel’s mean intensity across the time-course. Anatomical scans were first coregistered to the first EPI volume of the Food Pictures task and were then spatially normalized to Talairach space via an affine spatial transformation. Subject-level EPI data were only moved to standard space after subject-level regression analyses. All EPI data were left at the original spatial resolution (1.2×1.2×1.2mm^3^).

The EPI data collected during both tasks were separately analyzed at the subject-level using multiple linear regression models in AFNI’s 3dDeconvolve. For the FP task univariate analyses, the model included one regressor for each picture category (sweet, sour, salty, and objects). These regressors were constructed by convolution of a gamma-variate hemodynamic response function with a boxcar function having an 8-second width beginning at the onset of each presentation block. For the Taste Perception task univariate analyses, the model included one 16-second block regressor for each tastant type (sweet, sour, salty, and neutral) and one 4-second block regressor for wash/swallow events. The regression model also included regressors of non-interest to account for each run’s mean, linear, quadratic, and cubic signal trends, as well as the 6 normalized motion parameters (3 translations, 3 rotations) computed during the volume registration preprocessing.

We additionally generated subject-level regression coefficient maps for use in the multivariate ROI and searchlight analyses. For both tasks, we generated a new subject-level regression model, which modeled each run segment (8 total; see task design above) separately, so that all conditions of both tasks would have eight beta coefficient maps for the purposes of model training and testing.

## Analyses

### Imaging Analyses - Univariate

We generated statistical contrast maps at the group level to identify brain regions that exhibited shared activation for the sight of food pictures and the actual perception of taste. For this analysis, we used the subject-level univariate beta-coefficient maps to perform group-level random-effects analyses, using the AFNI program 3dttest++. For the Food Pictures task, we used the statistical contrast, all food pictures (sweet, sour, and salty) versus object pictures. For the Taste Perception task, we used the respective contrast, all tastants (sweet, sour, and salty) versus the neutral tastant. Both contrast maps were separately whole-volume corrected for multiple comparisons using a cluster-size FWE correction using non-parametric permutation tests (see Permutation testing section). We then performed a conjunction of the two independent contrast maps to identify brain voxels significantly activated by both tasks.

### Imaging Analyses - Multivariate

These analyses used a linear support-vector-machine (SVM) classification approach, implemented in The Decoding Toolbox(59), to classify tastants and food picture blocks based on their category labels. These SVM decoders were trained and tested on subject-level regression coefficients obtained from the Food Pictures and Taste Perception tasks, using leave-one-run-segment-out cross-validation. For this approach, we generated an independent set of ROIs from a previous study by our lab using the same gustatory imaging paradigm, at the same voxel resolution, but with a different group of subjects (12). The contrast used to produce these ROIs was the ‘all taste vs. neutral taste’ contrast (see Figure 3 from (12)), which generated six distinct insula clusters (Bilateral Mid-Insula, Bilateral Anterior Insula, Bilateral Ventral Anterior Insula; Figure 3). Within these ROIs, we compared the average pairwise classification accuracy vs. chance (50%) using one-sample signed permutation tests. This procedure generates an empirical distribution of parameter averages by randomly flipping the sign of individual parameter values within a sample 10,000 times. The p-value is the proportion of the empirical distribution above the average parameter (accuracy) value. These p-values were then FDR corrected for multiple comparisons. Main effects and interactions within these ROIs were tested with a permutation-based ANOVA, implemented in the *aovp* function in the R-library *lmPerm* (https://cran.r-project.org/web/packages/lmPerm/lmPerm.pdf).

The whole-volume MVPA searchlight analyses(60) allowed us to identify the average classification accuracy within a multivoxel searchlight, defined as a sphere with a four-voxel radius centered on each voxel in the brain (251 voxels/ 433 mm^3^ total). For every subject, we performed separate searchlight analyses for both imaging tasks. The outputs of these searchlight analyses were voxel-wise maps of average pairwise classification accuracy versus chance (50%). To evaluate the classification results at the group level, we warped the resulting classification maps to Talairach atlas space and applied a small amount of spatial smoothing (2.4 mm FWHM) to normalize the distribution of scores across the dataset. We then performed group-level random-effects analyses using the AFNI program 3dttest++ and applied a nonparametric permutation test to correct for multiple comparisons (see Permutation testing below for details). Through this procedure, we generated group-level classification accuracy maps for both the Food Pictures and Taste Perception tasks. We then created a conjunction of the two corrected classification maps, to identify shared brain regions present within both maps.

For the cross-modal decoding analyses, we trained the SVM decoder using the beta coefficients of the distinct tastes (sweet, sour, and salty) from the Taste Perception task and tested whether it could correctly predict the taste category of food picture blocks presented during the Food Pictures task, and vice versa. We performed this analysis within the insula ROIs described above, we corrected for multiple comparisons using a FDR correction. We also performed cross-modal decoding analyses using a multivariate searchlight approach, as described above.

### Pattern Similarity Analyses

We performed similarity analyses of the multivoxel patterns for tastes and food pictures within our mid-insula ROIs. For within modality analyses, we extracted the beta-coefficients (using AFNI’s 3dMaskdump) for all voxels within our insula ROIs, separately for odd and even runs, and then performed a voxel-wise correlation of odd and even runs for all task conditions (sweet, salty, sour). For between modality analyses, we performed a voxel-wise correlation of food picture and taste beta coefficients for all tastes (sweet, salty, and sour), using the full dataset beta coefficients. We Fisher-transformed the r-values and looked for an effect of *modality* (within vs. between) using a group level ANOVA.

### Voxel weight Analyses

We examined the distribution of voxel weights generated by the SVM model to classify tastes and pictures within our mid-insula ROIs. The SVM weights were generated in a transformation procedure described by (21), which allows for a SVM weights to be more clearly interpretable within the context of neuroimaging analyses. As with the pattern similarity analyses above, we performed split-half correlations of the voxel weights for within-modality analyses and correlations of the full dataset weights for between modality analyses, for each taste (sweet, salty, sour). We again Fisher-transformed the r-values and looked for an effect of *modality* (within vs. between) using a group level ANOVA.

### Object Pictures Analyses

To test whether images of non-foods could be distinguished in the same areas of the brain as images of foods, we ran a set of supplemental decoding analyses at the ROI and whole-brain level. For this analysis, we generated run-level beta coefficients at the participant level, using gamma-variate rather than block regressors, for a subset of the items presented in the non-food blocks of the Food Pictures task: lightbulbs, marbles, gloves, and innertubes. We subsequently ran multivariate decoding analyses on the object data within the insula ROIs and within a whole-brain searchlight, as we had for the food picture data, to determine the average accuracy for classifying these object pictures.

### Pleasantness Analyses

We performed a series of analyses to examine the effect of subjects’ self-reported pleasantness ratings for both food pictures and tastants on the activation during the respective imaging tasks. At the whole-volume level, we used separate amplitude-modulation regression analyses at the subject-level to identify whether the hemodynamic response to food pictures or tastants was modulated, at the trial-to-trial level, by participants self-reported ratings of the pleasantness of food pictures or tastes. In another approach, we used a linear-mixed-effects meta-analysis (using AFNI program 3dLME) to identify the variance explained at the group-level by participant’s pleasantness ratings both for tastants and food pictures. At the ROI level, we examined the effect of pleasantness ratings on the response to food pictures within the taste-responsive clusters of the insula, described above.

### Permutation testing for multiple-comparison correction

Multiple comparison correction was performed using AFNI’s 3dClustSim, within a whole-volume temporal signal-to-noise ratio (TSNR) mask. This mask was constructed from the intersection of the EPI scan windows for all subjects, for both tasks, (after transformation to Talairach space) with a brain mask in atlas space (Fig. 2c). The mask was then subjected to a TSNR threshold, such that all remaining voxels within the mask had an average un-smoothed TSNR of 10 or greater. For one-sample t-tests, this program will randomly flip the sign of individual datasets within a sample 10,000 times. This process generates an empirical distribution of cluster-size at the desired cluster-defining threshold (in this case, p < 0.001). The clusters which survive correction were those larger than 95% of the clusters within this empirical cluster-size distribution.

## Author Contributions

JAA, SJG, and AM designed research; JAA, AGL, and JEI performed research; JAA and AGL analyzed data; JAA, AGL, SJG, and AM wrote the paper.

## Acknowledgments

The authors would like to thank Sean Marrett, Martin Hebart, the NIMH Section on Instrumentation and the NIH Clinical Center pharmacy for their assistance with various aspects of the design and execution of this study. This study was supported by the Intramural Research Program of the National Institute of Mental Health, National Institutes of Health, and it was conducted under NIH Clinical Study Protocol 93-M-0170 (ZIA MH002920). Clinical-trials.gov ID: NCT00001360.

