## Supplemental Materials for "Tasting Pictures: Viewing Images of Foods Evokes Taste-Quality-Specific Activity in Gustatory Insular Cortex"

**This PDF file includes:**

Supplementary text
Figures S1 to S4
Tables S1 to S2

### Supplemental Methods

*Online Experiments*

We used Amazon Mechanical Turk (mTurK) to gather behavioral data from a large, diverse online sample. These online experiments allowed us to generate a novel set of food pictures to present to our subjects and to estimate certain inferred properties of the foods within these pictures, such as taste quality, nameability, and pleasantness.

*Foods with Characteristic Tastes*: For our first task, we prompted subjects to list foods that were characteristically sweet, salty, and sour. 105 mTurk workers participated in this Human Intelligence Task (HIT). Each worker wrote 5 unique foods that best represent sweet, sour, and salty foods. We took the top 6 responses generated for each taste category (ie. potato chips, bacon, pretzels, popcorn, french fries, and saltines for the salty taste category) and located a set of 28 unique pictures of each food using Google Images. We edited each food picture presented on a gray background.

*Identify Dominant Taste:* Having generated an image set of characteristically sweet, sour, and salty foods, we next wanted to ensure that subjects would reliably label these foods with the correct taste. We created a new task to measure the dominant taste of the image presented. We presented 452 workers with our food pictures, and they selected 'sweet', 'sour', 'salty', or 'not sure' per picture shown within a HIT. We used a catch-trial as well as reaction time (rt, rt >= lower quartile) to control for unreliable data. Mean correct label for final trial: salty = 96.0%, sour = 67.1%, and sweet = 98.0%. Through this process, we identified certain foods, such as sauerkraut and pickles, which had incorrect or inconsistent taste labeling. We added both new sour images as well as different sour foods to correct for these low label ratings.

*Quantifying Taste*: We next had subjects rate the tastes of these foods dimensionally, rather than categorically, as many foods contain a mixture of primary tastes, and may not serve as prime examples of one particular taste quality. We had 560 mTurk workers rate each depicted food on how sweet, sour, and salty it appeared to be. Within each HIT, workers rated one picture from each food type on three numerical scales which ranged from 0 (Not *sweet, sour, or salty* at all) to 5 (really *sweet, sour, or salty)*. A food naming task ('What is this picture?') was used to measure correct identification of these foods. Our final stimulus set was composed of foods that were highest in their respective taste category (sweet, salty, or sour) and low in the other two categories. Sweet foods: cake, donut, honey, ice cream; Sour foods: grapefruit, lemon, lemondrop/lemon candy, lime; Salty: (potato) chips, french fries, pretzels, saltines/crackers (Figure S1). In the final food picture stimuli set, mean naming accuracy = 93.6%. Mean taste ratings for each food stimulus type were plotted within a three-dimensional 'taste space' (Figure S1) and show a clear separation between the three food categories on the basis of their characteristic taste quality. A K-means clustering algorithm applied to the taste ratings from these food types created 3 distinct clusters: sweet, salty, and sour images, and showed 94.1% similarity as to what we predicted.

*Inferred Pleasantness:* We also asked online participants to rate the overall pleasantness of the foods within our picture set. 140 mTurk workers rated the final set of pictures on "how *pleasant* is the food in the image above?" They rated on a scale of 0 (not at all pleasant) to 7 (Really Pleasant). Results showed mean pleasantness for sweet = 4.9(1.5), sour = 4.2(1.5), salty = 4.3(1.5). We did observe an overall effect of pleasantness among the taste categories ($p < 0.001$), with pictures of sweet foods rated as overall more pleasant than salty or sour foods ($p < 0.001$) and no difference between pictures of salty and sour foods ($p = 0.07$).

*Familiarity of non-food objects:* We also generated a set of familiar non-food objects to serve as control pictures in our neuroimaging experiment. We selected foods which bore some resemblance to the items in our food picture set in terms of color, shape, and size. This set of objects consisted of: marbles, incandescent light bulbs, fluorescent light bulbs, pencils, tennis balls, basketballs, baseball gloves and innertubes. We located 14 unique pictures of each object (112 total) and asked workers on mTurk to rate those objects on their familiarity. workers rated 8 pictures per "HIT" on a scale of 0 (Not familiar) to 5 (Really familiar). 140 workers rated at least one picture from each category, resulting in 10 ratings per picture. We filtered data based on lower-quartile reaction times ( >= 6.25 seconds) and a control task of naming the object (must name object correctly). Object naming accuracy = 91.1% with a mean familiarity ratings = 4.42.

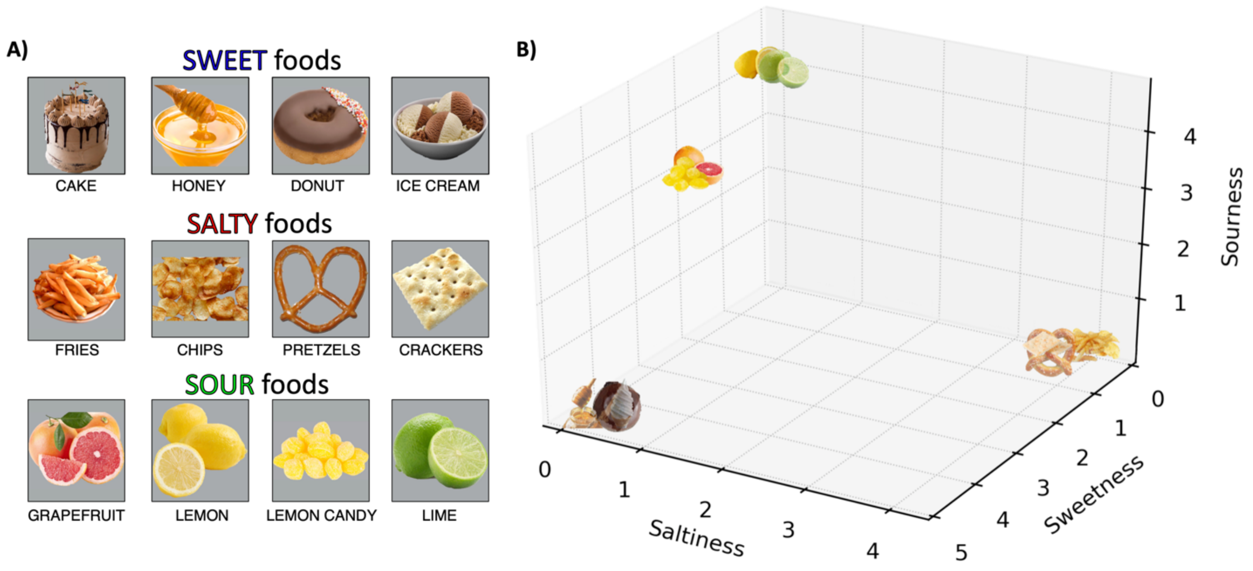


**Figure S1**: Characteristically sweet, sour, and salty foods selected and rated by an online sample of participants. Online participants were prompted to generate a list of characteristically sweet, sour, and salty foods. A) A set of food images was generated using that list and rated by another sample of online participants according to how characteristically sweet, salty, or sour those foods appeared to be. B) Our final stimulus set was composed of foods that were highest in their respective taste category (sweet, salty, or sour) and low in the other two categories.

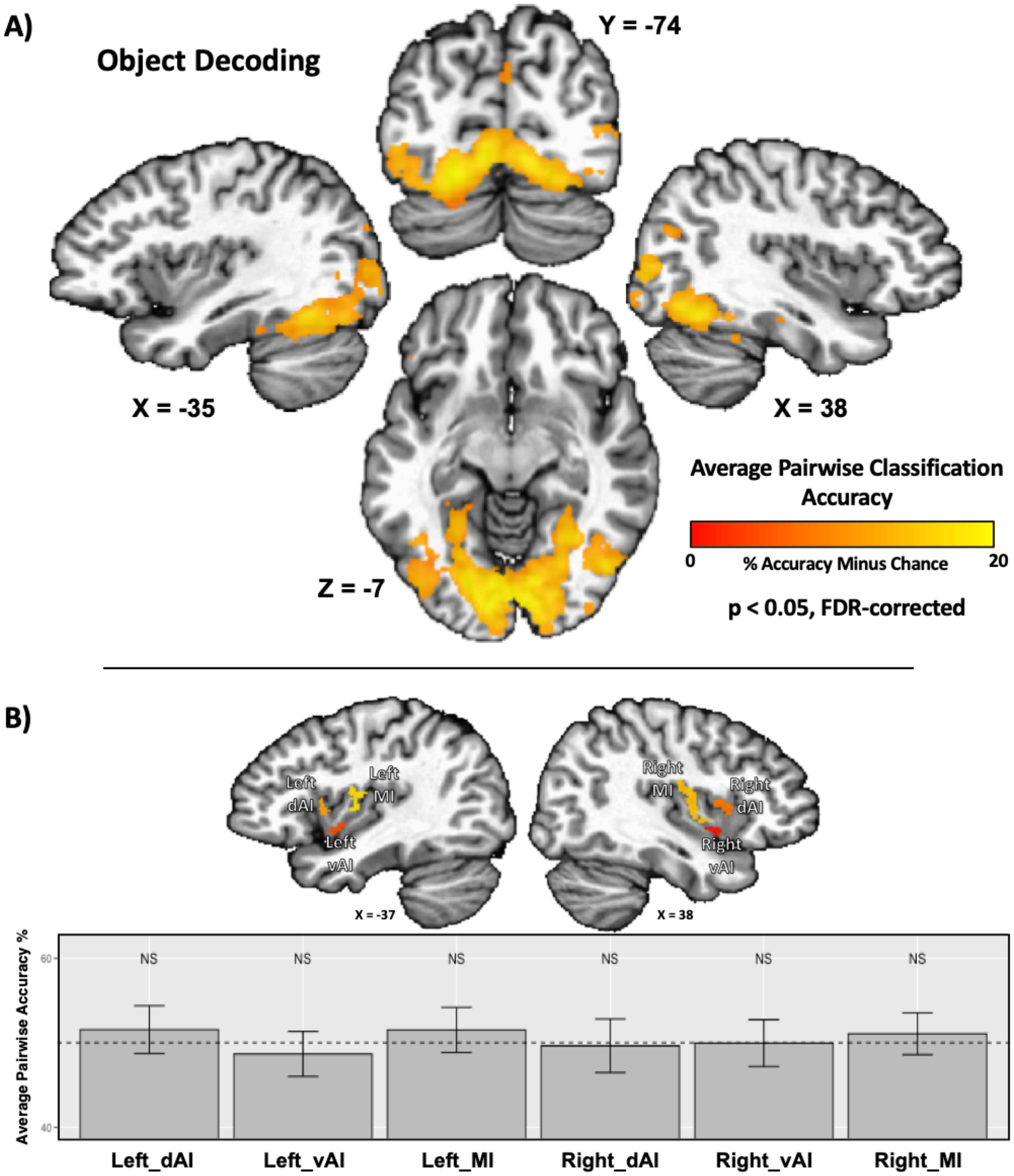


**Figure S2**: Object Decoding Analyses. A set of multivariate analyses were performed to decode pictures of non-food objects from the food pictures task. A) The searchlight analyses were able to reliably classify the object pictures within widespread areas of the occipitotemporal cortex, including primary visual cortex and much of the ventral visual processing stream, as was also seen when decoding food pictures. However, this set of regions was limited to occipitotemporal areas and did not include other areas able to decode food pictures, such as the orbitofrontal cortex, amygdala, or insula. B) ROI analyses within the insula confirm that classification accuracy for object pictures was no greater than chance.

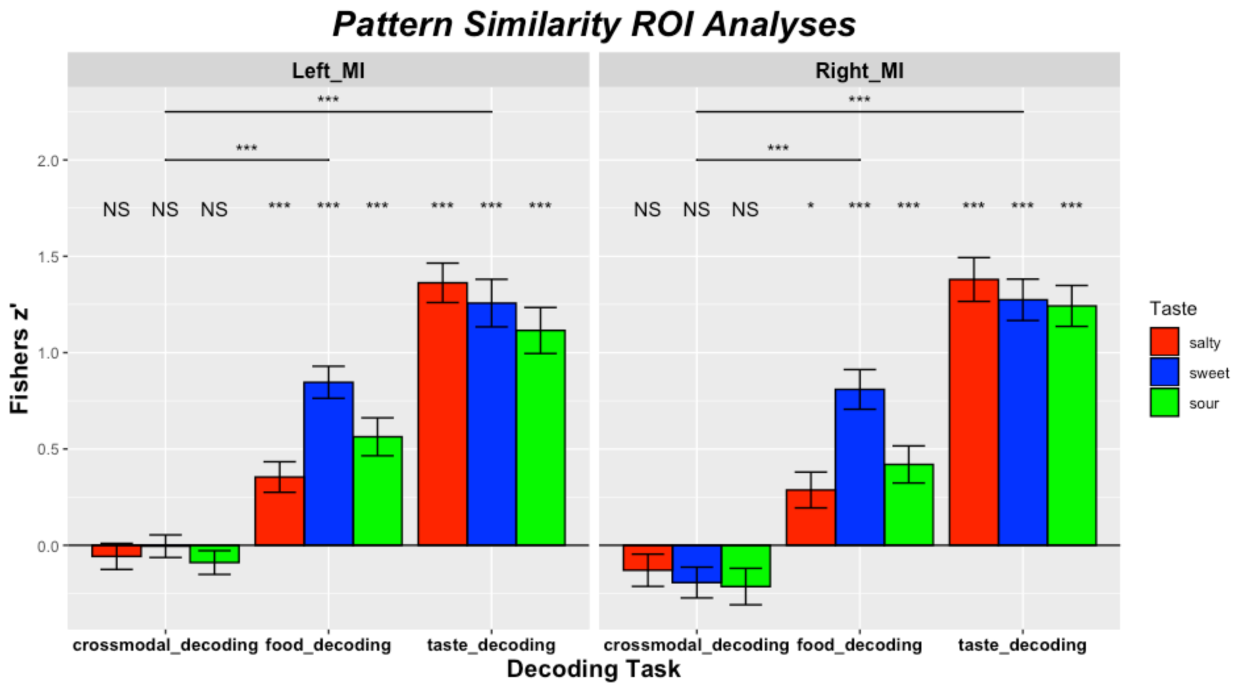


**Figure S3: Pattern Similarity Analyses**
We examined the similarity of multivariate patterns produced by tastes and food pictures within the mid-insula ROIs, both within and between modality. Using split-half correlations of food picture and taste data, we determined that the patterns produced by both of these tasks had high within-modal similarity for all taste conditions. When we examined the correlations between modality (crossmodal decoding), we observed that the similarity of all taste conditions was no different than zero and was significantly less than similarity within modality.

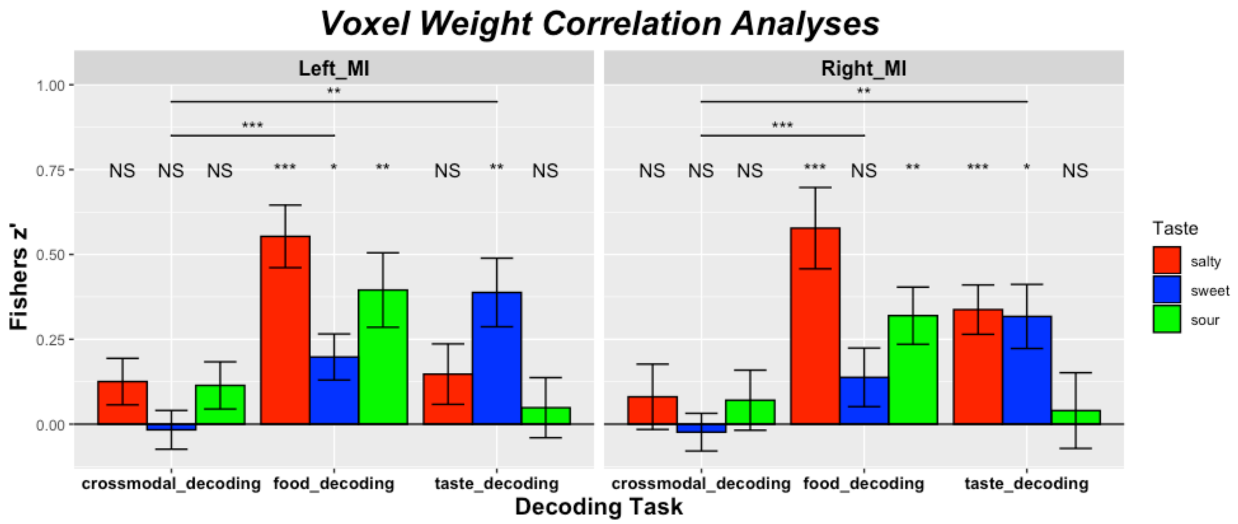


**Figure S4: Voxel Weight Correlation Analyses**
We also examined the distribution of voxel weights generated by our SVM models to separately classify tastes and food pictures within our mid-insula ROIs. These parameters indicate the voxels within each ROI which are most informative for predicting a particular taste or food picture category. We calculated the spatial correlation of these voxel weights on a subject-by-subject basis, both within modality (split-half) and between modality and examined the average correlations at the group level. Again, we observed significant within modality correlations, and the between modal correlations were no different than zero, as well as significantly less than within modal correlations.

**Table S1.** Behavioral Results

| | Pleasantness | Taste Contrast* | | | Intensity | Taste contrast* | | |
|---|---|---|---|---|---|---|---|---|
| Taste | Mean (SD) | Vs neutral | Vs salty | Vs sour | Mean (SD) | Vs neutral | Vs salty | Vs sour |
| Sweet | 4.41 (1.33) | **< 0.001** | **< 0.001** | **< 0.001** | 6.57 (1.92) | **< 0.001** | **0.001** | **< 0.001** |
| Sour | 3.18 (1.34) | 0.99 | **< 0.001** | | 5.13 (2.17) | **< 0.001** | .25 | |
| Salty | 2.08 (1.44) | **< 0.001** | | | 5.67 (2.00) | **< 0.001** | | |
| Neutral | 3.06 (0.80) | | | | 2.23 (2.23) | | | |

*listed p-values are the results of Bonferroni-corrected one-sample t-tests. Significant results are emphasized in bold font.

**Table S2**. Multivariate classification of food pictures and tastes within taste responsive regions of the insula

| ***Condition*** | **L dMI** | p* | **R dMI** | p* | **L dAI** | p* | **R dAI** | p* | **L vAI** | p* | **R vAI** | p* |
|---|---|---|---|---|---|---|---|---|---|---|---|---|
| Food Pictures | 71% | **<0.001** | 65% | **0.002** | 57% | 0.09 | 56% | 0.14 | 65% | **<0.001** | 62% | **0.002** |
| Taste | 63% | **0.002** | 67% | **<0.001** | 64% | **<0.001** | 65% | **<0.001** | 55% | 0.25 | 62% | **0.008** |
| Crossmodal Food → Taste | 52% | 0.54 | 50% | 0.81 | 49% | 0.81 | 50% | 0.81 | 53% | 0.11 | 50% | 0.81 |
| Crossmodal Taste → Food | 50% | 0.88 | 53% | 0.30 | 49% | 0.81 | 52% | 0.67 | 51% | 0.55 | 49% | 0.69 |

*listed p-values are the results of FDR-corrected one-sample permutation tests. Significant results are emphasized in bold font. dMI – Mid-Insula; vAI – Ventral Anterior Insula; dAI – Dorsal Anterior Insula
